# Coordinated circulating and tissue-based T cell responses precede xenograft rejection

**DOI:** 10.1101/2025.11.07.687259

**Authors:** Ekaterina Novikova, Elizabeth Severa, Han Chen, Elizabeth Doepke, Fiorella Chacon, Farshid Fathi, Nathan Suek, Benjamin Vermette, Robin Bombardi, Eloi Schmauch, Simon H. Williams, Jacqueline I. Kim, Imad Aljabban, Tal Eitan, Ian S. Jaffe, Grace Hammond, Karen Khalil, Aprajita Mattoo, Vasishta Tatapudi, Edward Skolnik, Elaina P. Weldon, David Ayares, Anoma Nellore, Megan Sykes, Adam D. Griesemer, Jeffrey M. Stern, Brendan J. Keating, Robert A. Montgomery, Ramin Sedaghat Herati

## Abstract

Despite the life-saving successes of solid organ transplantation, the number of individuals needing organ transplant far exceeds the number of organs available for use each year. Porcine xenotransplantation, or the use of pig organs for transplantation in people, holds substantial promise but xenograft rejection in humans is poorly understood. T cell rejection by the host immune system is a major challenge for human allografts and may limit the longevity of porcine xenografts. To study the xenograft rejection, we evaluated T cell responses and repertoire dynamics across tissues following porcine thymokidney transplantation in a decedent model over 61 days after bilateral native kidney nephrectomy. Despite induction with anti-thymocyte globulin and ongoing immune suppression consisting of rituximab, corticosteroids, calcineurin inhibition, and mycophenolate mofetil, human T cell infiltration of the xenograft was observed and was associated with xenograft dysfunction. Longitudinal analysis of T cell clonotypes in biopsies of thymokidney revealed accumulation of clonal human CD4 and CD8 T cell responses. Moreover, circulating activated T cells, including circulating T follicular helper (cTfh), were xeno-reactive and increased in frequency around rejection events. We confirmed clonal dominance of a single CD8 clonotype – identified as donor-reactive in a mixed lymphocyte reaction – in the circulation leading up to the acute cellular rejection event. Following re-treatment with anti-thymocyte globulin and intensification of corticosteroids, the T cell clonotypes were dramatically diminished in frequency in thymokidney and lymph nodes, though not eliminated. Over time of observation, the T cell clonotypes were shared across multiple compartments, including xenograft, circulation and lymph nodes and formed clonal families with known xeno-reactive clonotypes, suggesting a coordinated immune response against a limited pool of antigenic targets. Together, these data demonstrate T cell repertoire dynamics across tissues in the setting of xenograft rejection and highlight opportunities for early surveillance, prediction and potential intervention.

**One-sentence summary:** After pig-to-human kidney xenotransplantation, xenoreactive T cells form clonotypic families across blood, graft, and lymphoid tissues, indicating a coordinated and targeted response.

## Introduction

Kidney transplantation, the definitive renal replacement therapy, is used to treat tens of thousands of people annually in the United States. However, a critical shortage of compatible donor kidneys results in substantial morbidity and mortality (1). Recent advances in gene editing methods have enabled the knockout of xeno-antigen targets of antibodies mediating hyperacute rejection (2,3), thereby facilitating successful short-term transplantation of porcine kidneys (4,5) into brain-dead human decedents (6–10). A limited number of living individuals have received porcine organs, with graft survival reported for up to 6 months (11).

Despite the use of porcine genetic modification and multi-pronged immunosuppression, xenograft rejection is likely to remain a major impediment to long-term graft function due to the antigenic distance between humans and pigs (3). This genetic difference predisposes xenografts to acute rejection, which may involve both cellular and humoral mechanisms. Adaptive immune responses against the xenograft include T and B cell responses. Human CD4 and CD8 T cells can be activated through recognition of porcine antigens, either when presented on recipient APCs (indirect recognition (12,13)) or directly by SLA-peptide complexes on porcine APCs, though the latter is less common due to SLA/HLA incompatibility (14–16). In addition to acute cellular rejection, antibody-mediated rejection (AMR) can also compromise graft function either independently or concurrently with T cell infiltration. CD4 T cells can also contribute to AMR through activation of T follicular helper cells (Tfh) in germinal centers which has been observed in animal models for xenotransplantation (17) and may underlie some episodes of AMR. Furthermore, existence and contribution of porcine-reactive memory T cells to xenotransplantation is poorly understood but could be important for distinguishing whether the xenograft would be at high risk of rejection. Understanding the contribution of T cells to acute cellular rejection following renal xenotransplantation will be important to help advance this therapeutic modality.

The study of rejection has been limited in living allograft and xenograft recipients to date, due to the limited ability to perform repeated and invasive sampling which could lead to graft and patient compromise. Circulating immune cells can serve as a “window” into tissue immune events, provided that the mechanistic links between circulation and the tissue are understood. In a study of human kidney allografts, donor-reactive T cells expanded in circulation following transplantation. While these clonotypes were also detectable in rejecting grafts, the dominant graft-infiltrating clones were present, but not dominant in peripheral blood (18), underscoring the complexity of intercompartmental dynamics and need for deeper mechanistic insight to bridge these compartments. The clinical relevance is further emphasized by detection of donor-reactive clones contributing to kidney transplant rejection in unstimulated circulating pre-transplant repertoire (19). However, these links have not been established for xenotransplantation, particularly in the face of expanded immunosuppression regimens. Clonal overlaps between the kidney and lymph nodes have not been examined in both allo- and xenotransplantation studies in humans. The decedent model (6,10) overcomes many of these limitations by enabling longitudinal sampling across tissue compartments following xenotransplantation, providing a unique opportunity to define the mechanistic insight into T cell responses during rejection.

In a recent 61-day xenotransplantation study, a human decedent underwent bilateral native nephrectomy followed by transplantation of a pig kidney with thymus autograft (“thymokidney”) from an α-1,3-galactosyltransferase knockout (*GGTA1* KO) pig (20–23). We identified human T cells infiltrating the xenograft which was associated with xenograft dysfunction and showed that these included effector T cells identified as donor-reactive in pre-transplant mixed lymphocyte cultures (15,22,23). Longitudinal studies of T cell populations within xenograft biopsies confirmed the accumulation of specific, clonally expanded human CD4 and CD8 T cell responses directly within the organ. Moreover, analysis of the recipient’s blood revealed increased frequencies of circulating activated T cells (including cTfh) that were reactive against the xenograft, particularly around rejection events. We confirmed dominance of a single donor-reactive CD8 T cell clonotype in the blood immediately preceding acute cellular rejection. While retreatment with rATG and addition of intensified corticosteroids decreased the frequency of xeno-reactive T cells in the thymokidney, lymph nodes and blood, xeno-reactive clonotypes persisted. These findings illustrate the tissue-specific dynamics of the T cell repertoire during xenograft rejection, thereby highlighting potential avenues for earlier surveillance and intervention.

## Results

Porcine renal xenotransplantation was performed in a brain-dead individual (hereafter referred to as the decedent) following bilateral native nephrectomy (**Fig. 1A**), which was described in (6,22,23). Briefly, the decedent underwent induction with rabbit anti-thymocyte globulin (rATG), rituximab, high-dose corticosteroids, and plasmapheresis, followed by maintenance therapy with calcineurin inhibition, corticosteroids, and mycophenolate mofetil (**Fig. 1B**). Circulating T cells diminished following induction therapy, then started to recover around 3 weeks after rATG treatment (**Fig. S1A)**, whereas total leukocytes showed smaller, variable changes (**Fig. S1B**). Two episodes of rejection were identified based on decline of creatinine clearance (22). At post-operative day 33, renal biopsy demonstrated human IgA deposition in the glomeruli which was associated with reduced creatinine clearance. Plasma exchange and eculizumab were administered, resulting in normalisation of creatinine clearance to baseline levels and partial renal function recovery. However, renal function was impaired again starting on day 45, and xenograft biopsy identified acute cellular rejection on days 45 and 49. In response, rATG, belatacept, pegcetacoplan, and plasmapheresis were administered on days 49, 52 and 54. During the study period, regular sampling was performed for peripheral blood and thymokidney biopsies, and decedent iliac and porcine lymph nodes were recovered prior to transplantation and at the end of the study. The study concluded on day 61 per protocol.

**Figure 1.**
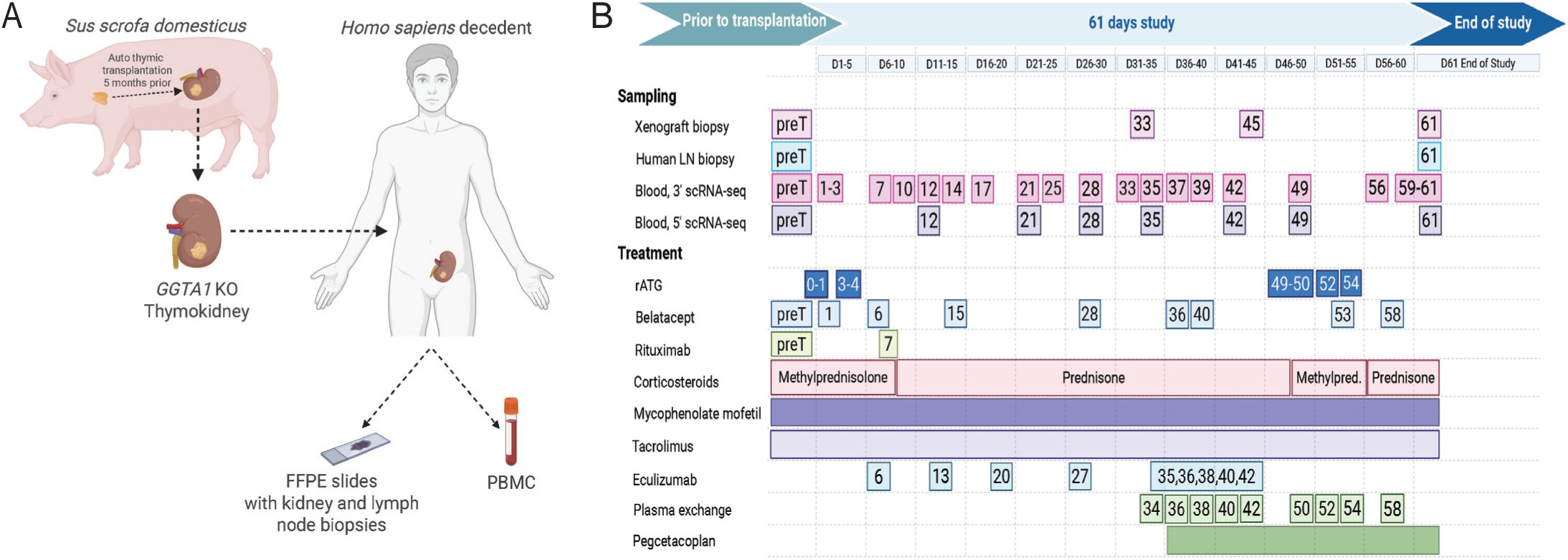
Overview of the 61-day *GGTA1* KO thymokidney transplantation study. **1A.** Schematic overview of the experimental setup, illustrating the transplantation of a *GGTA1* KO thymokidney from *Sus scrofa domesticus* into a human decedent. **1B**. Timeline detailing clinical events, sampling points, and immunosuppressive treatment regimen. LN – lymph node, rATG – rabbit antithymoglobulin, preT – pretransplantation time point

### Human T cells infiltrate thymokidney

To investigate the clonal dynamics of the human lymphocytic infiltrate, we performed bulk human antigen-receptor sequencing on mRNA recovered from formalin-fixed, paraffin-embedded (FFPE) tissue biopsy samples for *TCRα*, *TCRβ*, *TCRγ*, *TCRδ*, *IgH*, *IgL*, and *IgK* sequences (**Fig. 2A**). This approach did not recover human antigen-receptors in porcine xenograft prior to xenotransplant and only a few *TCRα* and no *TCRβ*, *TCRγ* and *TCRδ* in porcine lymph nodes, indicating little cross-reactivity for this assay for pig lymphocytes (**Fig. 2B, S2A, S2B**). Then, xenograft biopsies were evaluated on days 28, 33, 45, and at day 61. We quantified the number of human TCR clonotypes for the α, b, γ, and δ chains, defined as having a unique complementarity determining region (CDR)-3 nucleotide sequence (**Fig. 2B, S2B**). Across all time points, we identified few *TCRγ* or *TCRδ* receptor sequences in the xenograft (**Fig. S2B**). However, human *TCRα* and *TCRβ* receptor sequences and *IgH*, *IgK*, and IgL B cell receptor (BCR) sequences were identified in xenograft following transplantation and in the porcine and human lymph nodes at the end of study (**Fig. 2B, S2C-F**). These results establish the ability to profile human antigen receptors using FFPE biopsies of porcine xenografts.

**Figure 2.**
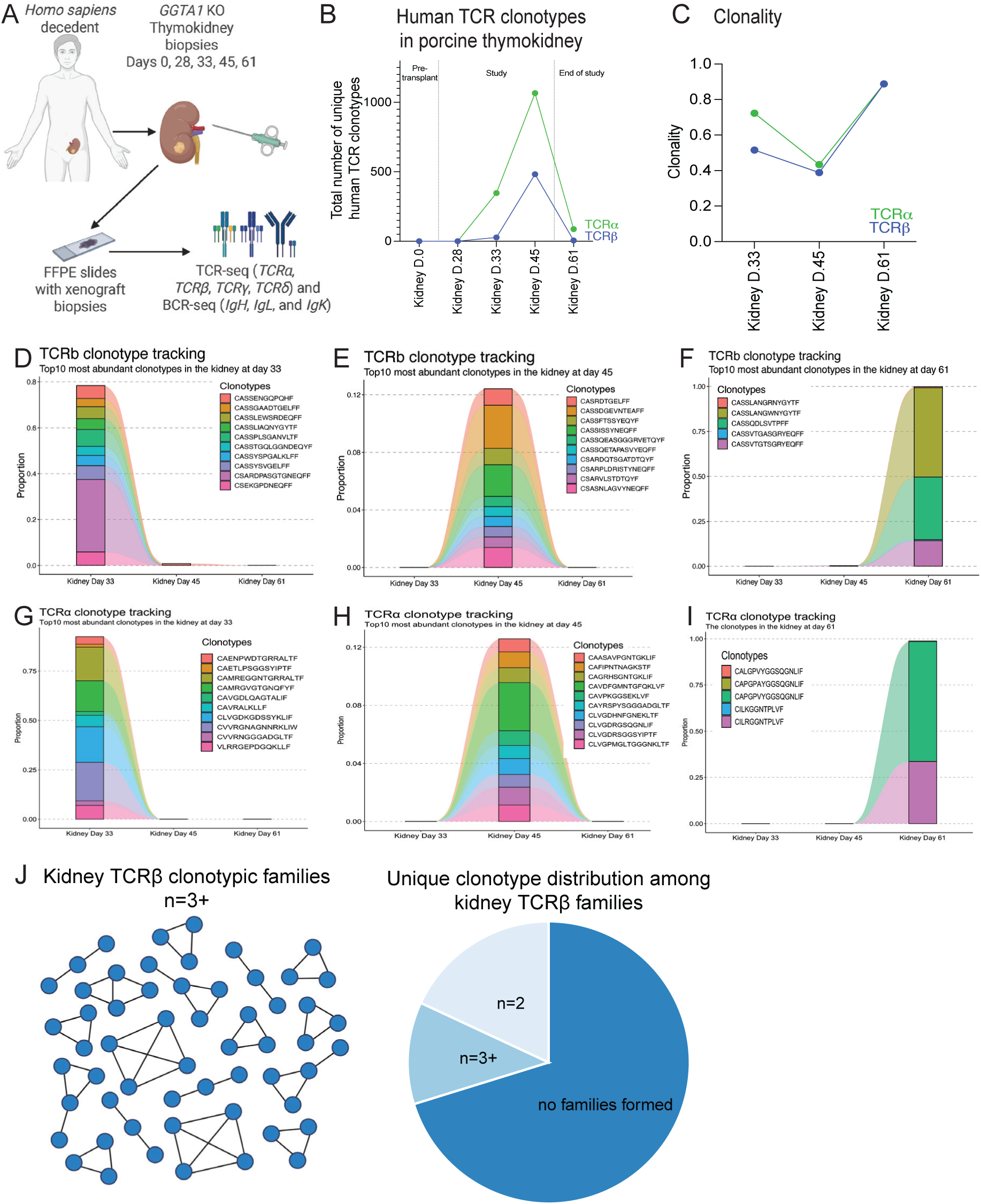
Human T cells infiltrate xenograft following transplantation. **2A.** Bulk TCR and BCR sequencing was conducted on RNA isolated from FFPE slides of thymokidney biopsies obtained prior to transplantation and at days 28, 33, 45 and 61 post-transplantation, as well as from a porcine lymph node biopsy and decedent iliac lymph nodes prior to transplantation and at the end of study. **2B.** Total number of unique human TCRα and b clonotypes, identified by unique CDR3 region in the xenograft over time. **2C.** Clonality of thymokidney samples for TCRα and TCRb over time. **2D**-**F.** Top 10 most abundant TCRb clonotypes identified in the thymokidney at each respective time point: day 33 (2D), day 45 (2E), and day 61 (2F), and tracking of their presence across other thymokidney time points. **2G-I**. TCRα clonotypes at day 33 (2H), 45 (2I), and 61 (2J), and tracking of their presence across other thymokidney time points. **2J.** Thymokidney TCRβ sequences form clonotypic families. Only families with 3 or more members are shown. Of the 487 unique clonotypes in the xenograft, 57 formed families with 3 or more members, 88 formed families with 2 members, and 342 did not form families.

We next asked how T cell infiltration changed over time. Though no *TCRα* and *TCRβ* cells were identified at day 28, *TCRα* and *TCRβ* cells were identified in the xenograft biopsies at day 33 and peaked at day 45 **(Fig. 2B**). The clonality dynamics for corresponsing timepoints are shown in **Fig. 2C**. Similarly, human B cells were detected as early as day 28 and peaked at day 45 **(Fig. S2E**). Repeated administration of rATG at day 49, however, resulted in >90% decline in *TCRα* and *TCRβ* in the xenograft by day 61 whereas iliac lymph nodes had *TCRα* and TCRβ (**Fig. S2C**). These data suggest rATG can indeed deplete xenograft-infiltrating αβT cells **(Fig. 2B**).

To further investigate T cell dynamics, we tracked the most abundant clonotypes in the xenograft at each time point to determine persistence over time, first focusing on *TCRβ* which exhibits more diversity than the *TCRα.* We identified the top 10 most abundant human *TCRβ* clonotypes in the xenograft biopsies at day 33 (**Fig. 2D**), day 45 (**Fig. 2E**), and day 61 (**Fig. 2F**) across time points and assessed clonotype sharing based on exact matching of the CDR3 nucleotide sequence. At day 33, the top 10 clonotypes accounted for nearly 80% of the repertoire. By day 45, TCR clonality decreased (**Fig. 2C**), with the top 10 clonotypes comprising just over 12% of the repertoire. By day 61, few *TCRβ* were found, with only five unique amino acid sequences detected. All time points had minimal to no overlap within xenograft. A similar lack of clonotypic sharing across time was observed for *TCRα* clonotypes (**Fig. 2G-I**). However, the lack of sharing could have been partially due to the low number of clonotypes recovered overall. We then asked whether clonotypic families could be identified, suggesting reactivity to a limited number of epitopes. Using the GLIPH2 algorithm (24), we indeed identified many clonotypic families from thymokidney biopsies across time points (**Fig. 2J**) despite the low preponderance of identical *TCRβ*. Together, these data suggest that T cells may be responding to common antigenic targets.

Similarly, BCR clonal dynamics were assessed using MiXCR (25). Many clonal families were identified among the BCR repertoire, with clonotype expression changing as the study progressed (**Fig. S2G**). Linear mixed model analysis revealed a slight increase in the number of mutations of these families over time (**Fig. S2H**), with 2 examples of individual BCR clonal trees appearing distinct and differing in mutation change over time (**Fig. S2I-J**). This suggests a shift in BCR repertoire complexity over time that may be reflective of germinal center activity in the context of xenotransplantation.

### Circulating T cells exhibit increased activation as rejection approaches

Given that reactive T cells migrate to the xenograft from secondary lymphoid organs via the bloodstream, we next studied circulating T cell responses (**Fig. 3A**). We hypothesized that following depletion, newly activated T cells responding to the targets in the xenograft would express proteins associated with recent activation, such as CD38 and HLA-DR, as has previously been observed. Indeed, the proportion of CD38+ HLA-DR+ activated T cells increased within the first 10 days post-transplantation, likely reflecting intensive T cell reconstitution following depletion (**Fig. 3B-C, S3A**) which has been observed following allotransplantation (26–28). Subsequently, the frequency of activated CD4+ and CD8+ T cells declined then began rising again, peaking at day 35. At the peak, 13% of CD4 T cells and 72% of CD8 T cells in circulation expressed CD38 and HLA-DR (**Fig. 3B-C**). Other phenotypic changes were observed in circulating T cells around day 35, with activated CD4⁺ and CD8⁺ T cells displaying increased CD38 expression (**Fig. 3D**). The high frequency of activated T cells in circulation persisted through day 49, which suggested that activated circulating T cells may serve as peripheral indicators of xenograft rejection.

**Figure 3.**
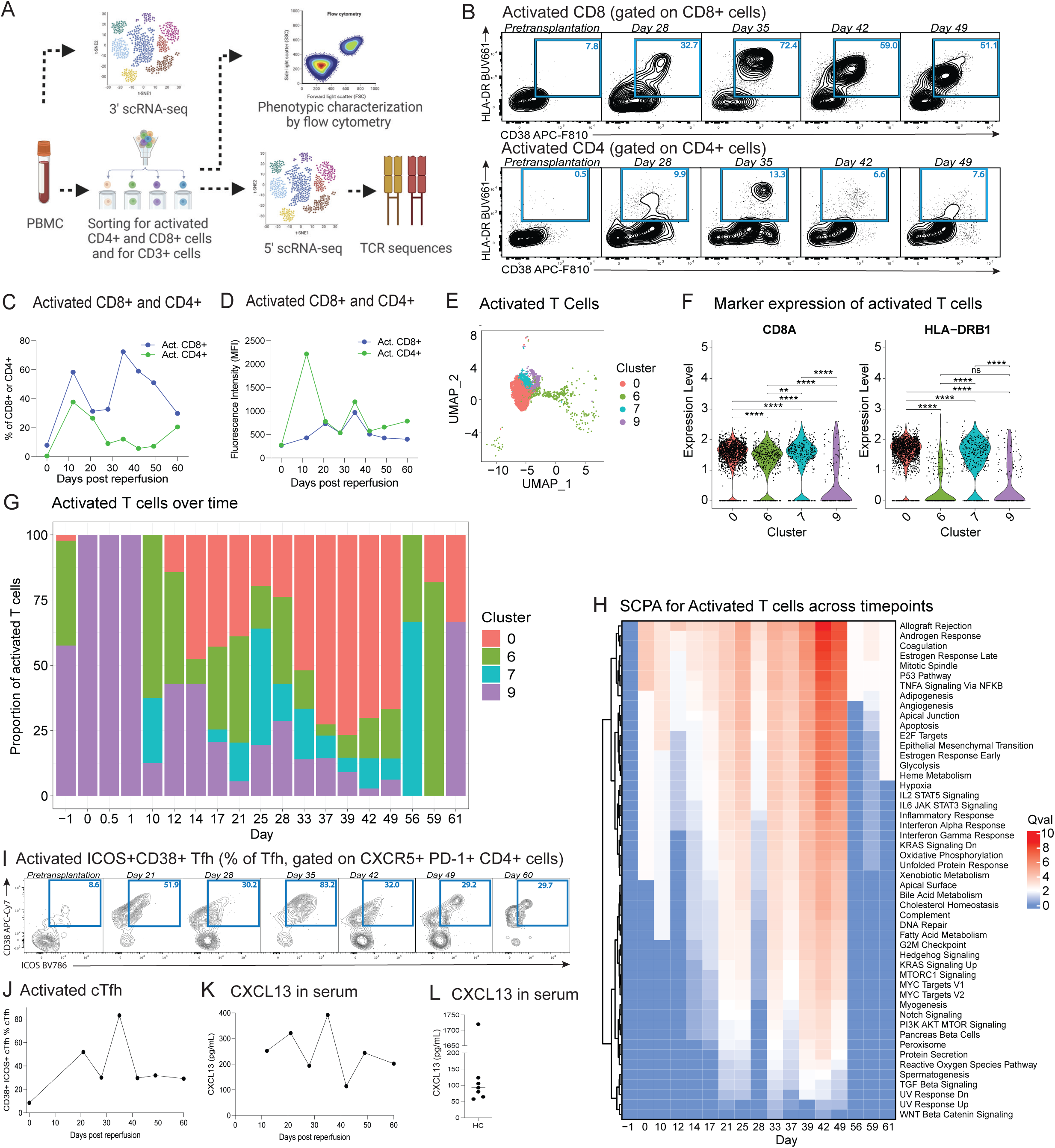
Circulating T cells exhibit increased activation as rejection approaches. **3A.** 3’-scRNAseq was performed on decedant PBMCs from prior to transplantation and from multiple time points after reperfusion. Additionally, PBMCs from selected time points were sorted to isolate activated CD4⁺ and CD8⁺ cells, and 5’ CITE-seq was performed on these sorted populations. **3B.** Activated CD8+ and CD4+ cells were identified based on expression of CD38 and HLA-DR. **3C.** The proportions of activated CD8+ and CD4+ cells, relative to their respective total CD8+ and CD4+ populations. **3D**. The Median Fluorescence Intensity of CD38 on CD8+ and CD4+ T cells. **3E.** UMAP of selected T cell clusters (0, 6, 7, and 9) from the 3’ scRNA-seq dataset from PBMC, representing activated T cells across all time points. **3F.** *CD8A* and *HLA-DRB1* expression across selected clusters confirm that the 3’ scRNA-seq subset accurately represents sorted T cell populations. Statistics were calculated via one-way ANOVA and post-hoc Tukey’s HSD test (**** p < 0.0001; *** p < 0.001; ** p < 0.01; * p < 0.05; ns p >= 0.05). **3G**. Proportion of activated T cells by cluster across different time points. **3H.** SCPA pathway expression for Hallmark genesets in activated T cells over time. **3I-3J.** Activated Tfh, identified by coexpression of ICOS and CD38. **3K.** CXCL13 level in serum of the long-term decedent over time. **3L.** CXCL13 level in serum of healthy individuals (HC).

To obtain a broader view of T cell repertoire dynamics over time following xenotransplantation, we evaluated T cells by single cell transcriptional profiling using 3’-profiling of unsorted PBMCs taken every 1-5 days following xenotransplantation. To identify activated T cells in this time-resolved dataset, we also generated 5’ CITE-seq dataset (29) from activated (CD38⁺ HLA-DR⁺) CD4⁺ and CD8⁺ T cells, as well as total CD3⁺ T cells, over multiple time points (days 0, 21, 28, 35, 42, 49, and 60) and used these sorted cell populations to robustly identify activated T cells in the unsorted data (**Fig. S3B**). Activated CD4+ and CD8+ T cells were identified through differential expression analysis and graph-based clustering. We identified differentially expressed genes between activated CD4⁺/CD8⁺ T cells and pre-transplant CD3⁺ T cells, generating a gene signature that was used to select highly activated clusters from the 3’ scRNA-seq dataset (**Fig. S3B**). Clusters 0, 6, 7, and 9 exhibited the highest alignment with this gene set and were selected for further analysis (**Fig. 3E-H, Fig. S3C-E**). The selection of clusters was further validated by elevated expression of CD8A and HLA-DRB1 (**Fig. 3F**), exhibiting similar marker profiles to what was seen and selected for in activated CD4⁺/CD8⁺ T cells.

Among these clusters, cluster 0 was dominant in the days around acute cellular rejection (**Fig. 3G, S3C**). This cluster was characterized by high grade expression of CD8A and HLA-DRB1 (**Fig. 3F**), as well as other genes associated with CD8+ T cell activation (**Fig. S3E**). Flow cytometry data also confirmed that the majority of activated CD3+ cells expressed CD8. The analysis of cluster dynamics (**Fig. 3G, Fig. S3C**) revealed that cluster 0 expanded following transplantation, increasing from day 33 onward and becoming the dominant population closer to rejection (days 33-49). In addition, cluster 0 had high levels of IFNg mRNA, with expression changing over time and corresponding with rejection (**Fig. S3D**). These findings support that the expansion of cluster 0 plays a role in the immune response leading up to rejection. The dynamics of this population are also supported by Single Cell Pathway Analysis (SCPA), with pathway expression fluctuating over time and in concordance with rejection events (**Fig. 3H**). Distinct pathway expression at days 33 and 49 suggests that these rejection events may differ in nature.

### T follicular helper cell signaling in circulation precedes the rejection

T follicular helper (Tfh) CD4+ T cells provide help to B cells in germinal centers. Although this is advantageous for long-lived humoral responses following vaccination, Tfh may also contribute to AMR by providing help to B cells. cTfh co-express CXCR5 and PD-1, and share phenotypic, functional, and transcriptional properties with lymphoid Tfh (30,31). To explore the role of Tfh in the rejection episodes observed, we first asked if cTfh proportion changed in circulation. We identified cTfh by flow cytometry (gating strategy shown in **Fig. S3A**, CXCR5 expression confirmed in **Fig. S3F**). At 2 weeks prior to observation of antibody deposition in the xenograft, the circulating frequency of cTfh rose dramatically (**Fig. S3G**), indicating the potential induction of germinal center activity. We then focused on the activated ICOS+ CD38+ cTfh subset which was previously identified to contain antigen-specific responding cells (30). Although the ICOS+ CD38+ cTfh subset was 8.6% pre-transplant, in line with our prior studies in healthy adults (32), this activated cTfh subset rose dramatically to 52% by day 21, slightly decreased to 30% at day 28, and peaking at 83% by day 35 post-transplant (**Fig. 3I-3J**). This dynamic pattern indicates initiation of germinal center activity around the time of antibody-mediated rejection.

To further explore the dynamics of cTfh responses, we sorted cTfh from PBMC from days 0, 21, 28, 35, 42, 49, and 60 and analyzed the transcriptional profiles using 5’ CITE-seq. Unsupervised clustering revealed three distinct Tfh clusters (**Fig. S3H**). The relative abundance of these clusters changed over time (**Fig. S3I**). Cluster 1 was dominant near day 35 whereas clusters 0 and 2 were dominant on days 28, 42 and 49 which preceded rejection events. Although clusters 0 and 2 had relatively higher expression of *ICOS* than cluster 1, cluster 1 had the highest expression of *HLA-DRB1* and *HLA-DRB5* genes (**Fig. S3J**). However, only a small number of cTfh were recovered per time point overall, thus investigation of cTfh subsets maybe warranted in future studies. Overall, however, these data support the notion that germinal center activity may have helped drive AMR and acute cellular rejection following xenotransplantation.

In addition, serum CXCL13 can serve as a circulating biomarker of GC initiation and correlate with lymphoid GC Tfh cell activity (33). Accordingly, we measured serum levels of CXCL13 at days 12, 21, 28, 35, 42, 49 and 60 post-transplant, as shown in **Fig. 3K**. Notably, CXCL13 levels were elevated at days 21 and 35 which coincinded with increases in activated cTfh at these time points (**Fig. 3J**), with all the time points higher than the median level of serum CXCL13 from healthy individuals (**Fig. 3L**). Put together, these data strongly implicate involvement of germinal centers and Tfh cell activity in the pathogenesis of xenotransplant rejection in this decedent.

### TCRβ clonotypes are shared among kidney, lymph nodes, and peripheral blood samples across multiple time points

We were next interested in understanding the clonotypic relationships underlying T cell activity across different time points. We used the TCR repertoire data from 5’ CITE-seq (**Fig. 3A**, bottom) on sorted activated (CD38⁺ HLA-DR⁺) CD4⁺ and CD8⁺ T cells (gating strategy in **Fig. S3A**), as well as total CD3⁺ T cells, from PBMC over multiple time points (days 0, 21, 28, 35, 42, 49, and 60). We first evaluated the top 10 most abundant TCRβ clonotypes at days 28, 35, 42 and 49 (**Fig. 4A**). Although multiple clonotypes were present across different time points in the blood, one clonotype (denoted CASSDGEVNTEAFF) was detected as early as day 28, expanded at days 35 and 42, and ultimately dominated the repertoire, comprising >60% of TCR sequences at day 49. Next, we asked whether the circulating CASSDGEVNTEAFF clonotype was also present in the xenograft biopsies or lymph nodes. We compared the TCR repertoire in the blood obtained from 5’ CITE-seq of sorted CD38+ HLA-DR+ CD4+ and CD8+ cells with the TCR identified in the xenograft biopsies and lymph nodes. The clonotype CASSDGEVNTEAFF was identified in the xenograft on day 45 (**Fig. 4B**), constituting approximately 2% of the TCRβ repertoire present at that time (**Fig. S4A**). The CASSDGEVNTEAFF clonotype was not present at detectable levels in the xenograft in other time points or lymph nodes following rATG and intensified corticosteroid treatment provided starting at day 49 when xenograft function partially recovered, indicating relevance of this clonotype in understanding xenograft rejection. We performed TCR sequencing on blood samples collected at early time points (days 5, 9, and 15) and detected CASSDGEVNTEAFF clonotype as early as day 9 (**Fig. S4B**), preceding T cell expansion in the blood and likely suggesting a memory phenotype and preexistence of this clonotype prior to transplantation.

**Figure 4.**
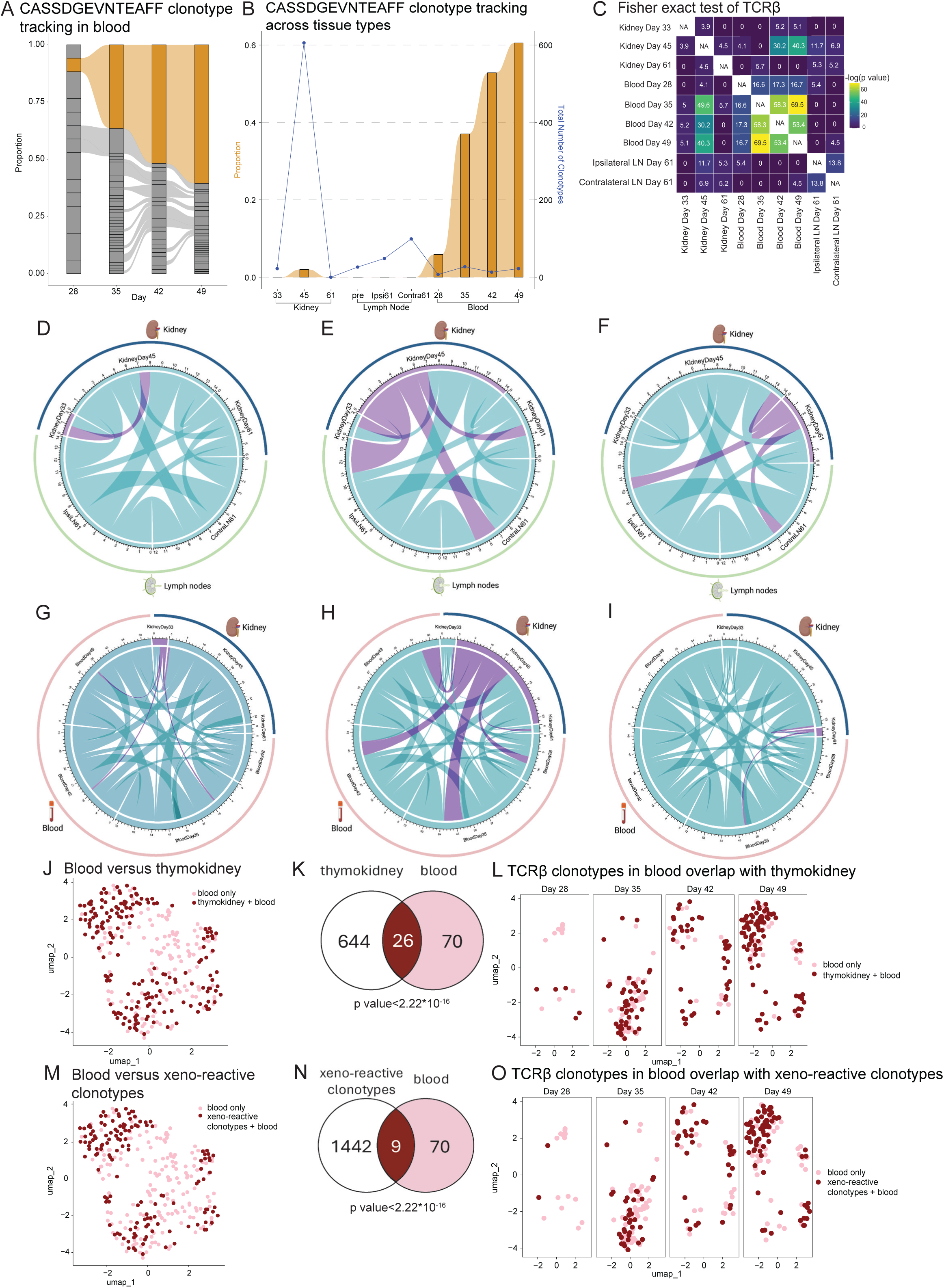
Clonal expansion of xenoreactive cells in xenograft and circulation. **4A.** Sharing of TCRβ in peripheral blood across days 28, 35, 42 and 49 post-transplant. Clonotype CASSDGEVNTEAFF shown in orange. **4B.** Tracking of TCRβ clonotype CASSDGEVNTEAFF across time and tissue compartments. The proportion of this clonotype is shown by the orange bars, and the total numbers of unique clonotypes per sample is shown by the blue line. **4C.** Heatmap of -log(p value) from Fisher’s Exact test of overlap of TCRβ clonotypes between xenograft, lymph node samples and PBMC. **4D.** Overlap of human TCRβ clonotypes in purple between xenograft at day 33 and clonotypes from other xenograft (Kidney) time points, as well as iliac ipsilateral (IpsiLN61) and contralateral (ContraLN61) lymph nodes at day 61. **4E.** Overlap of human TCRβ clonotypes between xenograft at day 45 and clonotypes from other xenograft time points and day 61 lymph nodes. **4F.** Overlap of TCRβ clonotypes between xenograft at day 61 and clonotypes from other xenograft time points and day 61 lymph nodes. **4G.** Overlap of TCRβ clonotypes between xenograft at day 33 and clonotypes from other xenograft time points, as well as blood samples at days 28, 35, 42 and 49. **4H.** Overlap of TCRβ clonotypes between xenograft at day 45 and clonotypes from other xenograft time points, as well as blood samples at days 28, 35, 42 and 49. **4I.** Overlap of TCRβ clonotypes between xenograft at day 61 and clonotypes from other xenograft time points, as well as blood samples at days 28, 35, 42 and 49. **4J.** UMAP of the shared TCRβ clonotypes present in peripheral blood and xenograft (dark red) and TCRβ clonotypes unique to blood (light pink). **4K**. TCRβ clonotypic sharing between blood and xenograft by number. **4L**. Shared TCRβ clonotypes present in peripheral blood and xenograft across time points. **4M.** UMAP of the shared TCRβ clonotypes present in peripheral blood and xeno-reactive clonotypes (dark red) and TCRβ clonotypes unique to blood (light pink) over time. **4N**. TCRβ clonotypic sharing between blood and xeno-reactive clonotypes by number. **4O**. Shared TCRβ clonotypes present in peripheral blood and xeno-reactive clonotypes (dark red) and TCRβ clonotypes unique to blood (light pink) over time.

We next considered clonotypic sharing across tissue compartments. In addition to the clonotype CASSDGEVNTEAFF, we also observed many other clonotypes shared across tissues (**Fig. 4C-I**). TCRβ clonotype sharing was limited between xenograft biopsy at day 33 and later xenograft biopsies (**Fig. 4D**). However, clonotypic sharing was prominent at day 45 with the day 61 xenograft biopsy as well as with the day 61 ipsilateral and contralateral inguinal lymph nodes (**Fig. 4E**). Following rATG and intensification of corticosteroid treatment, clonotypic sharing was also observed with the contemporaneous lymph node samples ( **Fig. 4F**), suggesting coordinated T cell responses across these compartments. Notably, rATG reduced the total number of clonotypes but did not completely eliminate T lymphocytes from the kidney or lymph nodes. To quantify TCRβ clonotype overlaps between different kidney and human lymph node samples, we performed Fisher’s exact test. The resulting p-values are represented as a heatmap of -log(p value) in **Fig. 4C**.

When examining TCRβ sharing between kidney and blood samples (**Fig. 4G-I**), we observed that the kidney shared only a few clonotypes with the blood at day 33 (**Fig. 4G**). However, at day 45 (**Fig. 4H**), the number of shared clonotypes increased, with clonotype exchange observed across all kidney and blood time points. By day 61 (**Fig. 4I**), only a single clonotype was shared between the kidney and blood, likely reflecting the impact of rATG administered at days 49, 50, 52, and 54, which may have more effectively depleted circulating T cells compared to those residing in the kidney and lymph nodes. Fisher exact test, shown as a -log(p value) in **Fig. 4C**, supports the TCRβ clonotype sharings between kidney and blood time points, with more overlaps observed closer to rejection. On a single-cell level, the overlaps between TCRβ clonotypes in blood and kidney (**Fig. 4J-K**) were found to change over time (**Fig. 4L**). Several clonotypes were shared between kidney and blood, highlighted in dark red in **Fig. 4J**, with 26 distinct clonotypes overlapping, mapping to 173 cells in the blood (**Fig. 4K**). This overlap shifts as time progresses, and is shown more prominently after rejection time points (**Fig. 4L**).

We have previously documented the marked expansion of donor-reactive clonotypes identified through recipient anti-donor pig mixed lymphocyte reaction using high-throughput TCRβ CDR3 sequencing of DNA from T circulating cells at various post-transplant timepoints (15,22,23). T cell clones infiltrating the rejecting kidney xenograft at POD 33 included donor-reactive clones with an effector phenotype. We interrogated the clonotypes in our current study for these xenoreactive clones to determine the kinetics of their expansion and entry into the xenograft in more detail. The clonotypes were shared between TCRβ blood and xeno-reactive clonotypes datasets (**Fig. 4M-O**). Nine unique clonotypes overlapped, which mapped to 140 cells in the blood (**Fig. 4N**). The overlap changes over time and appears more prominently closer to the rejection time points (**Fig. 4O**).

### TCRβ clonotypes form clonotypic families across multiple time points

Additionally, grouping TCRs by sequence has allowed identification of families that are likely reactive towards a shared antigen (24,34,35). Using the GLIPH2 algorithm, we screened our datasets to identify such clonal families. TCRβ clonotypes from lymph nodes at the end of study, known xenogeneic donor-reactive clonotypes (15,22,23), kidney-derived clonotypes, circulating clonotypes from activated CD4+/CD8+ and Tfh populations indeed formed clonotypic families (**Fig. 5A**). Interestingly, CASSDGEVNTEAFF also formed a clonotypic family with known xenogeneic donor-reactive clonotypes, suggesting its xeno-reactive nature.

**Figure 5.**
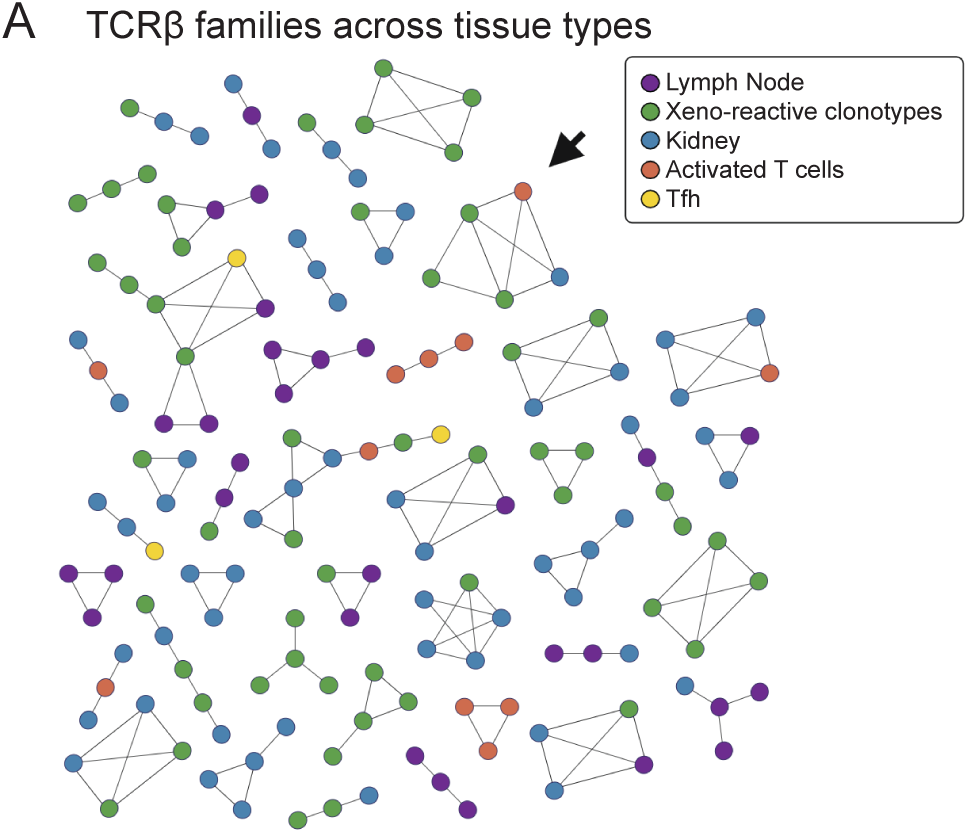
TCRβ clonotypes form clonotypic families across compartments. **5A.** GLIPH2-based clonal families form across peripheral blood, lymph node, kidney, Tfh, and xenogeneic donor-reactive clonotypes. Arrow indicates clonotype CASSDGEVNTEAFF.

Notably, circulating Tfh clonotypes were members of clonotypic families that included clonotypes from the thymokidney biopsies and clonotypes of proliferating T cells after mixed lymphocyte reaction using porcine cells. These findings suggest that these Tfh cells are likely xeno-reactive and might have contributed to AMR, although direct analysis of germinal center responses and antibody specificity would be needed to better understand this relationship.

Although the T cell repertoires in circulation and tissue compartments differ in size and cellular composition, the overlaps between compartments suggest a coordinated immune response following porcine renal xenotransplantation, in spite of immune suppression. Moreover, the peripheral blood may serve as a window into immune-mediated rejection occurring after solid organ transplantation.

## Discussion

Porcine renal xenotransplantation is a promising approach for renal replacement therapy and may help alleviate the organ shortage, but xenograft survival may be limited by rejection involving anti-xenograft human T cell responses. In this study, two episodes of xenograft rejection that impaired xenograft function were identified, necessitating escalation of immunosuppressive therapy. In the xenograft, human immune cells dynamics were evaluated through antigen receptor sequences, revealing a coordinated immune response. Circulating activated T cells, particularly CD8+ T cells and ICOS+CD38+ cTfh cells, increased in frequency leading up to rejection events. Although oligoclonal CD8+ T cell responses were observed in the xenograft, a single clonotype dominated in the circulation. Overall, xeno-rejection was associated with observation of shared clonotypes across tissue compartments and across time points. Sampling across compartments allowed richer characterization of xenograft rejection than was possible using any one modality individually.

Solid organ rejection has been challenging to study due to the need for coordinated sampling across different tissues, which is further limited by the uncertainty of the timing of onset of rejection. Although circulating T cell responses have been repeatedly evaluated (18,36), cross-compartment analysis in humans has been limited due to the inherent challenges of tissue sampling. This has been achieved in intestinal allograft recipients, in which serial surveillance biopsies are the standard screening method for rejection and such studies have provided unique understanding of the dynamic interplay between alloreactive T cells and lymphocyte dynamics between the circulation and the graft itself (37,38). The decedent model affords the opportunity of longitudinal sampling of tissues in the settings of kidney transplantation including repeated evaluation of the xenograft itself, which would otherwise not be reasonable in a living kidney recipient. In our results, we observed that recipient T cell activity preceded the clinical identification of rejection at day 33 and increased steadily until day 49 when immunosuppression was intensified. The circulating T cell repertoire was clonally linked to the xenograft and contained numerous xeno-reactive clonotypes. Notably, one clonotype (CASSDGEVNTEAFF) was markedly expanded, comprising more than 60% of the circulating repertoire by day 49. This largely expanded clonotype was also identified prior to xenotransplantation in the same individual and shown to be donor-reactive (15,22,23). Better understanding of T cell clonal dynamics after xenotransplantation will facilitate earlier detection of rejection, thereby allowing more timely intervention. Moreover, the dominance of a single clonotype (CASSDGEVNTEAFF) in circulating activated T cells but not in the xenograft suggests proliferation of these T cells outside of the xenograft, likely in lymphoid tissues, similar to what has been observed in allograft rejection in animal models (39,40). Future studies will need to directly sample lymphoid tissues in the midst of the xenograft rejection to understand the contribution of T cell trafficking to xenograft rejection. Distinguishing which responses involve the direct, indirect, and semi-direct pathways may also have implications for chronic rejection (41).

We observed indirect evidence of germinal center activity in the secondary lymphoid tissues. The dynamic expansion of circulating Tfh cells around the time of antibody-mediated rejection, clonal link between circulating Tfh, xenograft biopsy clonotypes and xeno-reactive clonotypes as well as increase in CXCL13 in serum suggests the initiation of germinal center responses, supporting the hypothesis that germinal center responses may contribute, at least in part, to AMR. Although lymphoid tissue was not sampled during the rejection in this study, future studies incorporating longitudinal lymphoid tissue sampling will be essential to directly establish the role of germinal centers in the rejection. Moreover, if the link between activated Tfh cells and germinal center activity is confirmed in the context of xenotransplantation across multiple studies, circulating activated Tfh cells could potentially serve as a predictive biomarker for antibody-mediated rejection.

The interspecies differences present in the current study likely led to rejection in spite of immunosuppression. A recent study comparing *Sus scrofa* and *Homo sapiens* genomes observed that over 80% of genes are one-to-one orthologs (42). Thus, substantial antigenic differences of the proteomes across these species likely remain as potential targets of the immune responses. However, we found oligoclonal TCRβ responses that formed clonotypic families that were present in the blood, lymph node, and xenograft. One possibility is that T cell rejection may have been preferentially directed against a limited set of xeno-antigens, in spite of the antigenic dissimilarity, perhaps due to dominance of memory T cell responses such as from prior food exposure (43) or because of restricted antigenic presentation. Another possibility is that there might have been T cell repertoire depletion due to conditioning regimens, or that the immunosuppressive regimen raised the threshold for T cell activation due to calcineurin inhibition that biased T cell responses. Moreover, identification of the specific antigens driving xenograft rejection may thus provide opportunities for improvement of the suitability of porcine organs through targeted mutations.

Some caveats apply. As a case study, generalizability may be limited. Future studies involving additional decedents will be needed to allow generalization of the observations, and impending Phase I clinical trials may provide another opportunity for evaluation of T cell responses. Additional efforts will be needed to understand the porcine antigens driving xeno-reactive T cell responses and the thresholds for T cell activation given complex immunosuppressive regimens. Although our study evaluated multiple tissues during the monitoring period, sampling of secondary lymphoid organs, including across time points, will be essential to understanding T cell immunity, particularly for indirect responses. Finally, extending the follow-up period beyond 60 days will allow greater understanding of whether chronic rejection will impair graft survival after porcine xenotransplantation.

Understanding the immune mechanisms underlying acute xenograft rejection is critical for prolonging the xenograft survival and advancing xenotransplantation. This study leverages the decedent model to perform multisite immune profiling across blood, lymph node and xenograft compartments, enabling a level of resolution rarely achievable in living recipients. Together, our findings reveal a coordinated and multicompartmental humoral and cellular response, including clonal T cell expansion and evidence of germinal center activity. These insights lay the groundwork for the development of biomarkers for early detection of both cellular and humoral rejection and improve understanding of optimal timing for immunosuppressive therapy. Moving forward, identifying the specific xenoantigens that drive rejection and directly sampling lymphoid tissues during rejection will be essential to inform targeted immune modulation and improve xenograft outcomes.

## Materials and methods

### Experimental model and study design

We used blood and tissue samples collected from a previously-described human decedent model of xenotransplantation (6,8–10,22). In this model, bilateral native nephrectomies were performed prior to the implantation of a single thymokidney graft, which had been procured from a *GGTA1* KO pig five months after thymic autotransplantation. All procedures were conducted under the oversight of the New York University Research on Decedents Oversight Committee (RDOC Protocol #22-001), in accordance with established protocols for research on human decedents. Induction and maintenance immunosuppression followed the FDA-approved standard of care for transplant patients. The study concluded on day 61 per protocol.

### Sample selection

Fresh PBMCs were isolated using the SepMate protocol (STEMCELL Technologies), followed by red blood cell lysis and cryopreservation with FBS+10% DMSO, at baseline (pre-transplant) and on days 1, 2, 3, 7, 10, 12, 14, 17, 21, 25, 28, 33, 35, 37, 39, 42, 49, 56, 59, 60, and 61 post-transplantation. Formalin-fixed paraffin-embedded (FFPE) slides containing core needle biopsies were obtained from 10 distinct sampling events from the decedent model, including the pre-transplant human iliac lymph node, pre-transplant pig lymph node, pre-transplant pig thymokidney, thymokidney at days 28, 33, 45, and 61 (end-of-study), perigraft porcine lymph node at day 61, human iliac ipsilateral and contralateral lymph nodes at day 61. Two 10 μm sections were collected per sample for subsequent antigen receptor profiling. Serum samples were collected from prior to transplantation and at days 12, 21, 28, 35, 42, 49, 60.

### High-throughput TCR sequencing in tissue biopsy samples

After RNA isolation from FFPE slides (the isolated RNA concentrations are shown in Supplemental Table 2), next-generation sequencing (NGS) libraries covering all TCR chains (*TCRα*, *TCRβ*, *TCRγ*, and *TCRδ*) and BCR chains (*IgH*, *IgL* and *IgK*) were generated using the iR-RepSeq+ service provided by iRepertoire, Inc. (Huntsville, AL, USA). The sequencing protocol used primer pairs for each V-D-J combination and incorporated dam-PCR technology, enabling the amplification of TCR chains in a single assay, incorporating unique molecular identifiers (UMIs) during the reverse transcription (RT) step to distinguish individual RNA molecules and reduce the impact of PCR duplicates and sequencing errors. The raw sequencing data were analyzed using R (version 4.3.2) and the library Immunarch (v. 1.0.0) (44). For TCR clonotype overlap analysis, unique clonotypes were identified based on their unique CDR3 amino acid sequences.

### 3’ scRNAseq

The cryopreserved PBMC across 22 time points were thawed in four batches and then loaded for standard 10X Genomics single-cell RNA-seq (v3.1 Chemistry, 10x Genomics, CA, USA).

### Flow cytometry and cell sorting

Cryopreserved PBMC from days 0, 21, 28, 35, 42, 49, and 60 were thawed and treated with Fc receptor blockade using Human TruStain FcX (BioLegend) and NovaBlock (Thermo Fisher Scientific), both at 1:100 dilution, for 10 min at room temperature. This was followed by surface antibody staining at room temperature for 30 min in the dark (Supplemental Table 2). For sample multiplexing, cells were stained with hashtag oligonucleotides for 30 min at room temperature (BioLegend). Acquisition and sorting were performed on a BD FACSymphony S6 Cell sorter using the 100 μm nozzle. Sorting gates included CD3+ T cells, activated CD4+ and CD8+ T cells (defined as CD38+ HLA-DR+) and CXCR5+ PD-1+ Tfh cells. The full gating strategy is shown in Figure S3A. Sorted cells were processed for 5’ CITE-seq using the Chromium Next GEM Single Cell 5′ HT Kit v2 (10x Genomics). Cells were pooled, loaded onto Chromium GEM-X Chips, and run on a Chromium controller (10x Genomics). Gene expression, TCR V(D)J, and surface protein expression libraries were produced using the 5′ Feature Barcode Kit, the Chromium Single Cell V(D)J Amplification Kit, and the Chromium Next GEM Single Cell 5′ Library Kit (10x Genomics), following the manufacturer’s instructions. Libraries were pooled and sequenced using the NovaSeq 6000 (Illumina).

### 5’ CITE-seq data processing, statistical analysis, and visualization

*Cellranger software v8.0.1 (10X Genomics) was used to align FASTQ files to the human genome (GRCh38), evaluate gene and protein expression, and exclude empty droplets via the Barcode Rank Plot. Seurat v4.3.0 was used for single cell library processing and assay integration. HTO demultiplexing* was conducted using CLR normalization (via *NormalizeData()*) and *HTODemux()*. Cells were filtered and excluded based on high mitochondrial counts, and low and high UMI counts. Additional processing included normalizing and scaling. For doublet detection, *scDblFinder()* was used, with an estimated doublet rate of 0.4% per 1000 cells. *HTODemux()* negatives were removed. scRepertoire v1.11.0 and Immunarch v0.9.1 was used for TCR and BCR sequence processing. Seurat allowed for integration, via SCTransform for normalization, *SelectIntegrationFeatures()* (utilizing 5000 features), and *FindIntegrationAnchors()* (utilizing the first 20 dimensions, rpca reduction, and 20 k.anchors). Azimuth v0.4.6 was utilized to aid in celltype assignment. SCPA v1.6.2 was used for pathway analysis and enrichment evaluation. All statistical analyses were performed using two-sided testing at α=0.05, including for one-way ANOVA and post-hoc Tukey’s HSD test in R as indicated. Turbogliph v0.99.2 was used for clonotypic relationships. Mixcr v4.3.2 and LMM lme4 v1.1-36 were used for BCR clonal families. Study schematics and figures were made using BioRender and Adobe Illustrator 2025 v29.2.1.

### Extracting transcriptional markers for CD38+ HLA-DR+ CD4⁺ and CD8⁺ T cells from the 3′ scRNA-seq dataset

Data was converted from .h5ad format to an alternative format (matrix.mtx.gz, barcodes.tsv.gz, features.tsv.gz) in Python 3.12.7 using scanpy v.1.10.3 and scipy 1.14.1. Processing and quality control evaluation was conducted as described above. Data were filtered to T cells via Azimuth assignment. Differentially-expressed genes between sorted activated CD4+ and CD8+ T cells and sorted pre-transplant CD3 cells were determined. Genes upregulated in activated CD4+ and CD8+ T cells were retained as a geneset and used to recluster T cells via PCA, where clusters 0, 6, 7, and 9 aligned most closely with the geneset and were extracted for further analysis.

### Serum CXCL13

Serum samples collected at 8 time points over the course of transplantation were thawed and transferred to a serum separator tubes (SST). Samples were allowed to clot for 30 minutes at room temperature before centrifugation for 15 minutes at 1000 x g. CXCL13 concentrations were measured using SimplePlex Human CXCL13/BLC/BCA-1 assay kit (Ella, ProteinSimple). Serum samples were diluted 2x with Sample Diluent and loaded into the cartridge according to the manufacturer’s instructions. Concentrations were reported in pg/mL.

## Data availability statement

5’ CITE-seq data were deposited into the Gene Expression Omnibus database under accession number GSE297896 (reviewer token: yncngumebjsrxaf), and are available at the following URL: https://www.ncbi.nlm.nih.gov/geo/query/acc.cgi?acc=GSE297896.

## Code availability statement

All scripts will be publicly available on Zenodo. Requests for resources or further information should be directed to the lead contact at NYU Grossman School of Medicine.

## Acknowledgements

We sincerely thank the families of the decedent for their generous donation to science. We thank the NYU Langone Health Nursing Leadership, NYU Transplant Research Team, and the NYU Langone Health Center for Biospecimen Research and Development (CBRD), Histology and Immunohistochemistry Laboratory (RRID:SCR_018304), NYU Surgical Intensive Care Unit Advanced Practice Providers, NYU Surgical Intensive Care Unit Nursing Staff, NYU Grossman School of Medicine’s Research on Decedent Oversight Committee, NYU Langone, Donor Care Unit, and LiveOn NY. We thank United Therapeutics for preparation of *GGTA1*-KO thymokidney. We thank Dr. Alexandre Loupy and his team for helpful discussions for data analysis and interpretation. We thank Dr. Massimo Mangiola for the assistance with the antibody crossmatch assays, which were instrumental in identifying the decedent. We are also grateful to Dr. Philip Sommer for the help with the decedent’s clinical care, to Dr. Sapna Mehta, clinical director of NYU Langone Transplant Institution, and to Dr. Ming Wu for the pathological evaluation of the xenograft biopsies.

## Contributions

Concept or study design: E.N., S.H.W., A.N., M.S., A.D.G., L.B., J.M.S., B.J.K., R.A.M., R.S.H.

Acquisition of data: E.N., H.C., E.D., F.C., F.F., N.S., B.V., J.I.K., I.A., T.E., I.S.A., G.H., K.K., E.P.W.

Analysis: E.N., E.S.^1^, H.C., R.B., E.S.^5,6^, A.N.

Manuscript preparation: E.N., E.S.^1^, R.S.H.

Decedent’s clinical care: A.M., V.T., E.S.^1,9^

Generation of GGTA1 knockout pig donors: D.A.

---

**Supplemental Figure 1.**
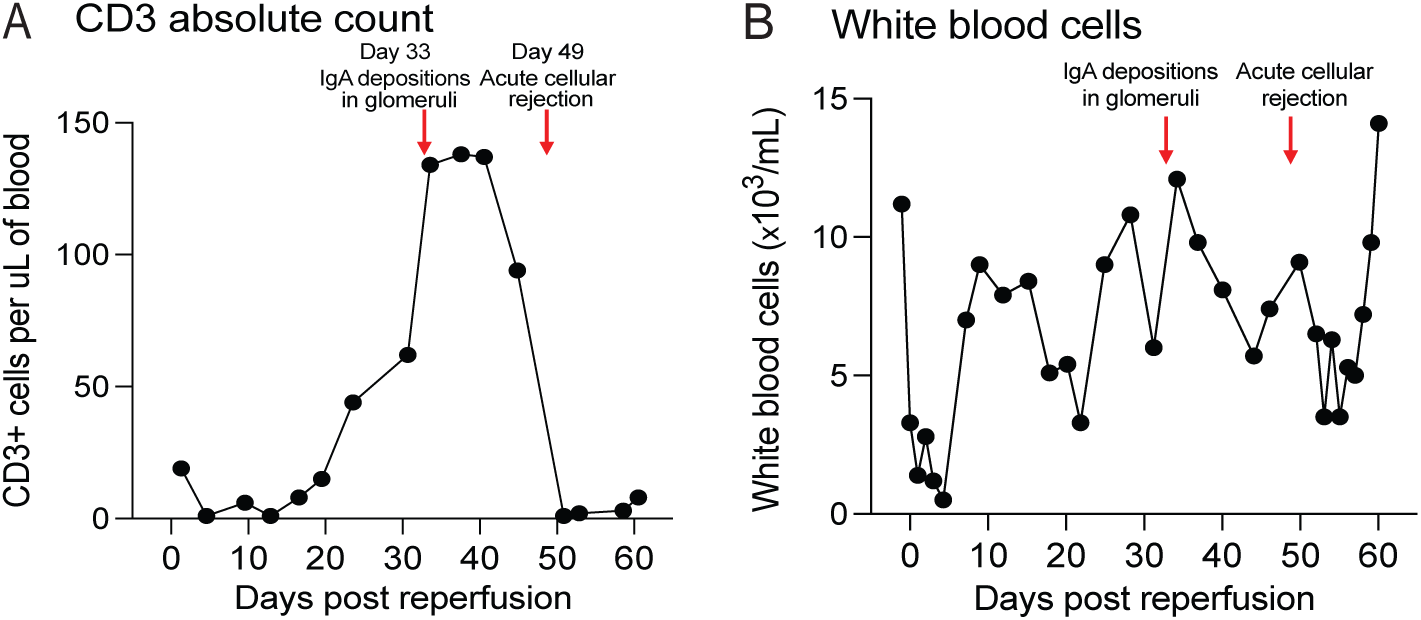
**S1A.** CD3 absolute count (cells/μL) over time. **S1B.** White blood cell (WBC) count (×10^3^/mL) over time.

**Supplemental Figure 2.**
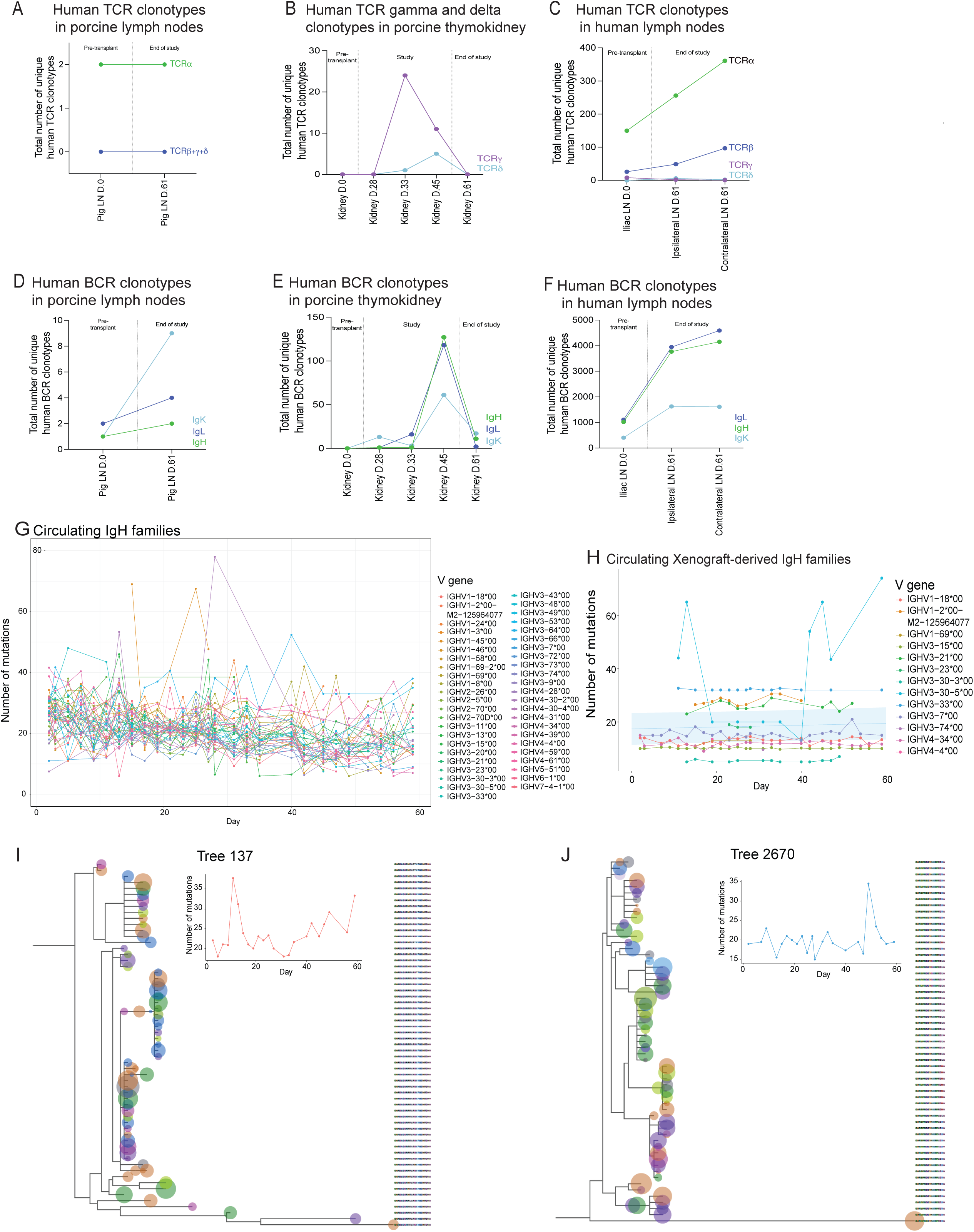
**S2A.** Total number of unique human TCRα (green) and TCRb, γ and δ (blue) clonotypes, identified by unique CDR3 region, in the porcine perigraft lymph nodes prior to transplantation and at the end of study. **S2B.** Total number of unique human TCR γ (purple) and δ (light blue) clonotypes, identified by unique CDR3 region, in the xenograft over time. **S2C.** Total number of unique human TCRα (green), TCRb (blue), γ (purple) and δ (light blue) clonotypes, identified by unique CDR3 region, in the human iliac lymph nodes prior to transplantation and at the end of study. **S2D.** Total number of unique human BCR clonotypes: IgH (green), IgL (blue) and IgK (light blue) in the porcine perigraft lymph nodes prior to transplantation and at the end of study. **S2E.** Total number of unique human BCR clonotypes: IgH (green), IgL (blue) and IgK (light blue) in the xenograft over time. **S2F.** Total number of unique human BCR clonotypes: IgH (green), IgL (blue) and IgK (light blue) in the human iliac lymph nodes prior to transplantation and at the end of study. **S2G.** BCR clonotype expression changes over time. Over 3000 clonal families (trees) were determined, and these were isolated to only IGH sequences overlapping with BCRs in the kidney (via iRepertoire), and displayed here grouped by V gene. **S2H.** Linear mixed model analysis suggests a slight increase in complexity over time, grouped by V gene. **S2I-J.** BCR clonal trees are distinct from each other and differ in mutation change over time. Mutation changes in tree 137 and 2670 are shown, alongside their phylogenetic trees.

**Supplemental Figure 3.**
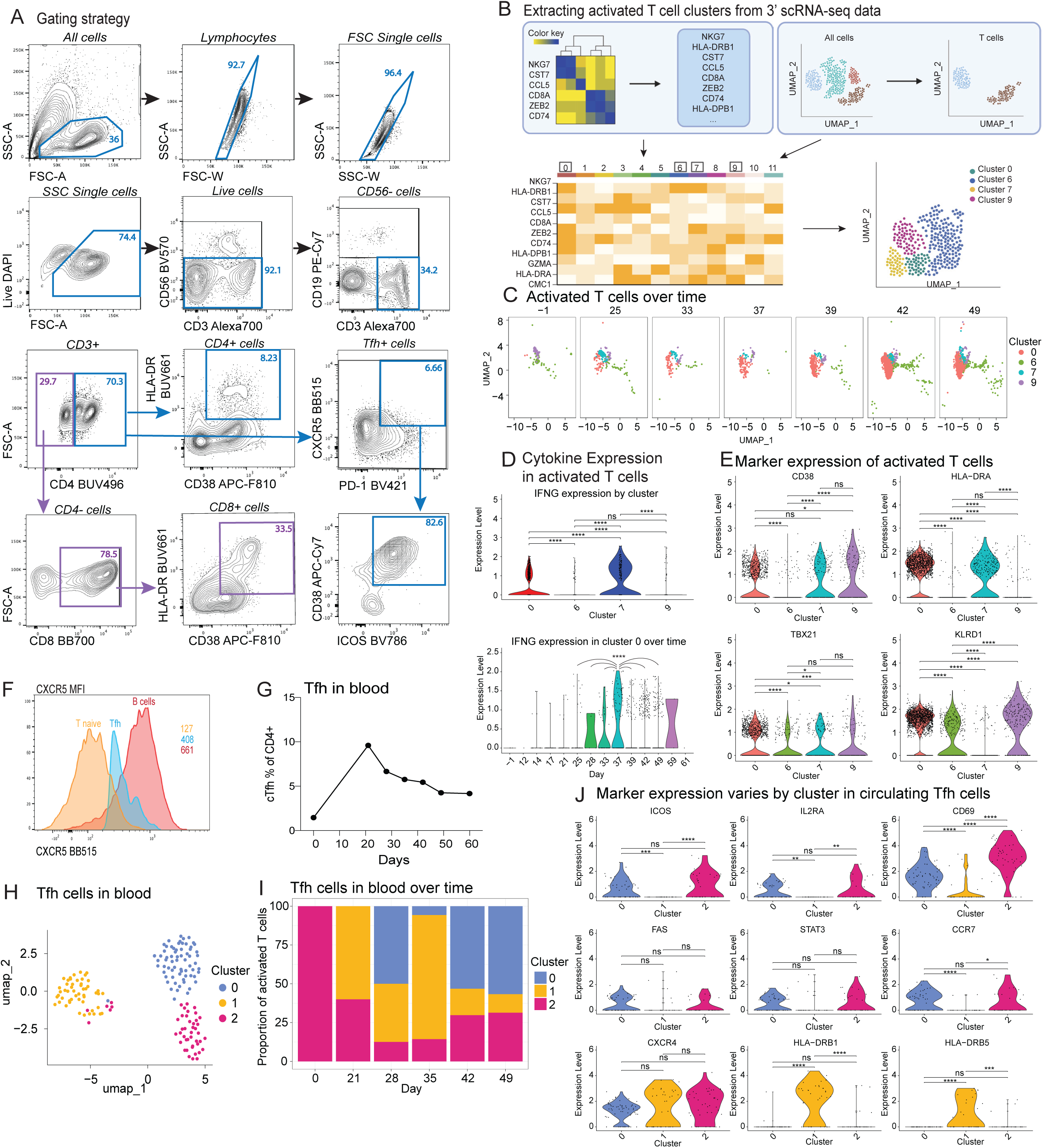
**S3A.**The full gating strategy to sort for CD3+ T cells, activated (CD38+ HLA-DR+) CD4+ and CD8+ cells, as well as CXCR5+ PD-1+ Tfh cells **S3B.** Extracting activation-associated transcriptional markers for CD4⁺ and CD8⁺ T cells from the 3′ scRNA-seq dataset. **(top)** Determine differentially-expressed genes between the activated CD4+ and CD8+ dataset and the pre-transplant CD3 dataset; keep markers upregulated in activated CD4+ and CD8+ dataset only. Use azimuth labels (l1) to filter 3’ scRNAseq dataset to just T cells **(bottom left)** Use markers from (top) to recluster 3’ scRNAseq T cells via PCA; clusters 0, 6, 7, and 9 align most closely with (top) markers. **(bottom right)** Cells of cluster 0, 6, 7, and 9 extracted for analysis **S3C.** Clusters of activated CD4+ and CD8+ over time; complement to 3G. Change in cluster composition over time. **S3D.** Cytokine gene expression for *IFNG* by cluster and over time for cluster 0. Statistics were calculated via one-way ANOVA and post-hoc Tukey’s HSD test (**** p < 0.0001; *** p < 0.001; ** p < 0.01; * p < 0.05; ns p >= 0.05). **S3E.** Expression of *CD38*, *HLA-DRA*, *TBX21* and *KLRD1* by scRNAseq. Statistics were calculated via one-way ANOVA and post-hoc Tukey’s HSD test (**** p < 0.0001; *** p < 0.001; ** p < 0.01; * p < 0.05; ns p >= 0.05). **S3F.** CXCR5 Median Fluorescence Intensity (MFI) for naive T cells (orange), T follicular helper cells (blue) and B cells (red). The values represent the MFI for each respective population: naive T cells (orange), Tfh cells (blue), and B cells (red). **S3G**. The frequency of circulating Tfh cells, identified as CXCR5+ PD-1+ CD4+ T cells **S3H.** UMAP of sorted Tfh cells in blood. **S3I.** Tfh clusters change contribution over time. **S3J.** Tfh clusters feature distinct expression profiles. Statistics were calculated via one-way ANOVA and post-hoc Tukey’s HSD test (**** p < 0.0001; *** p < 0.001; ** p < 0.01; * p < 0.05; ns p >= 0.05).

**Supplemental Figure 4.**
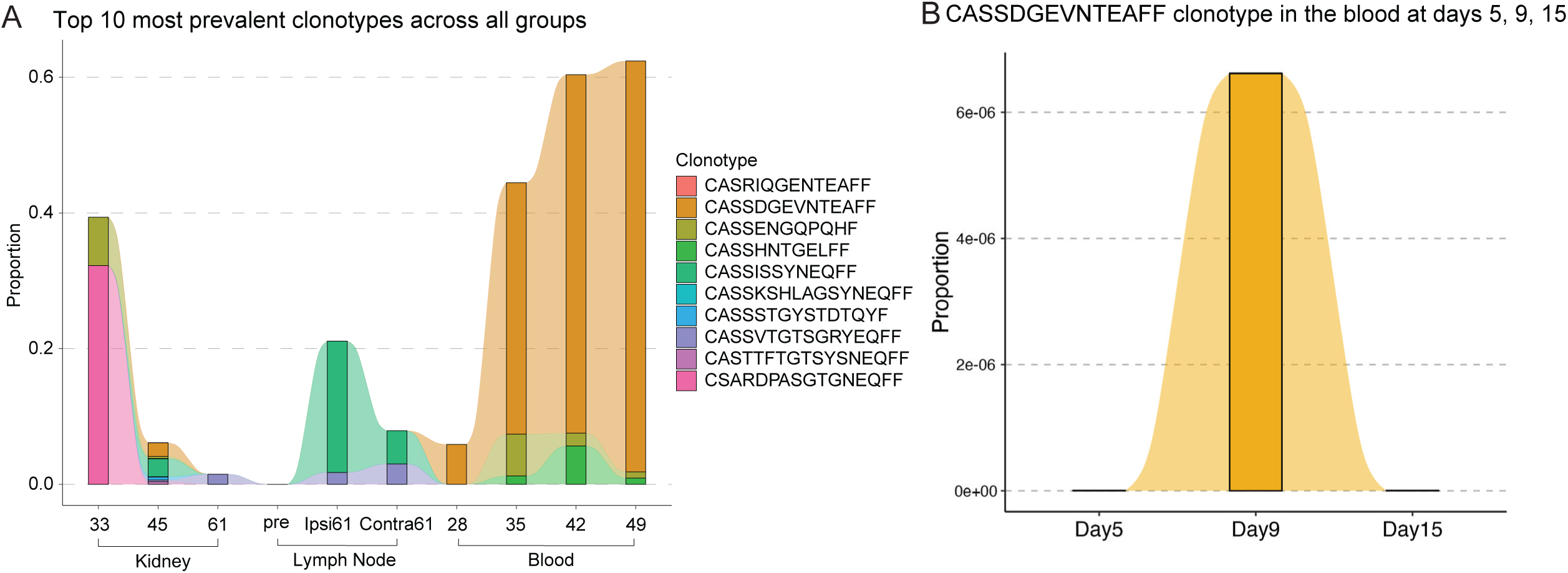
**S4A.** Top 10 most prevalent clonotypes across time and compartments. **S4B.** CASSDGEVNTEAFF clonotype tracking in the blood at early time points: day 5, day 9, day 15.

