## Supplemental figures and tables for "Coordinated circulating and tissue-based T cell responses precede xenograft rejection"

Supplemental Figure 1

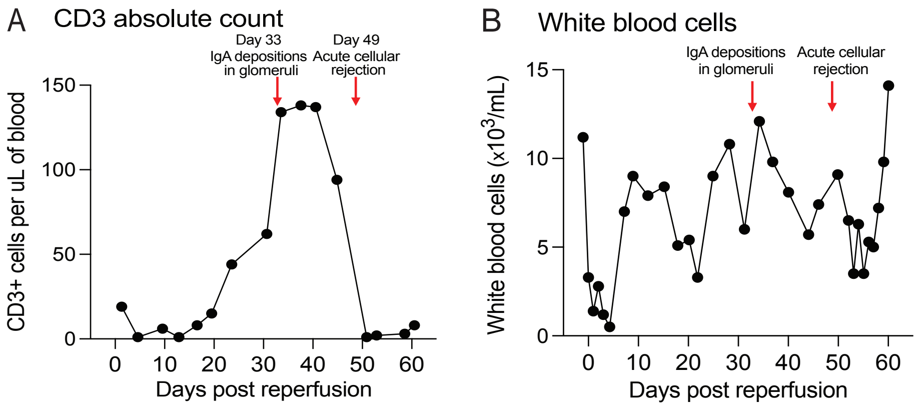

Supplemental Figure 2

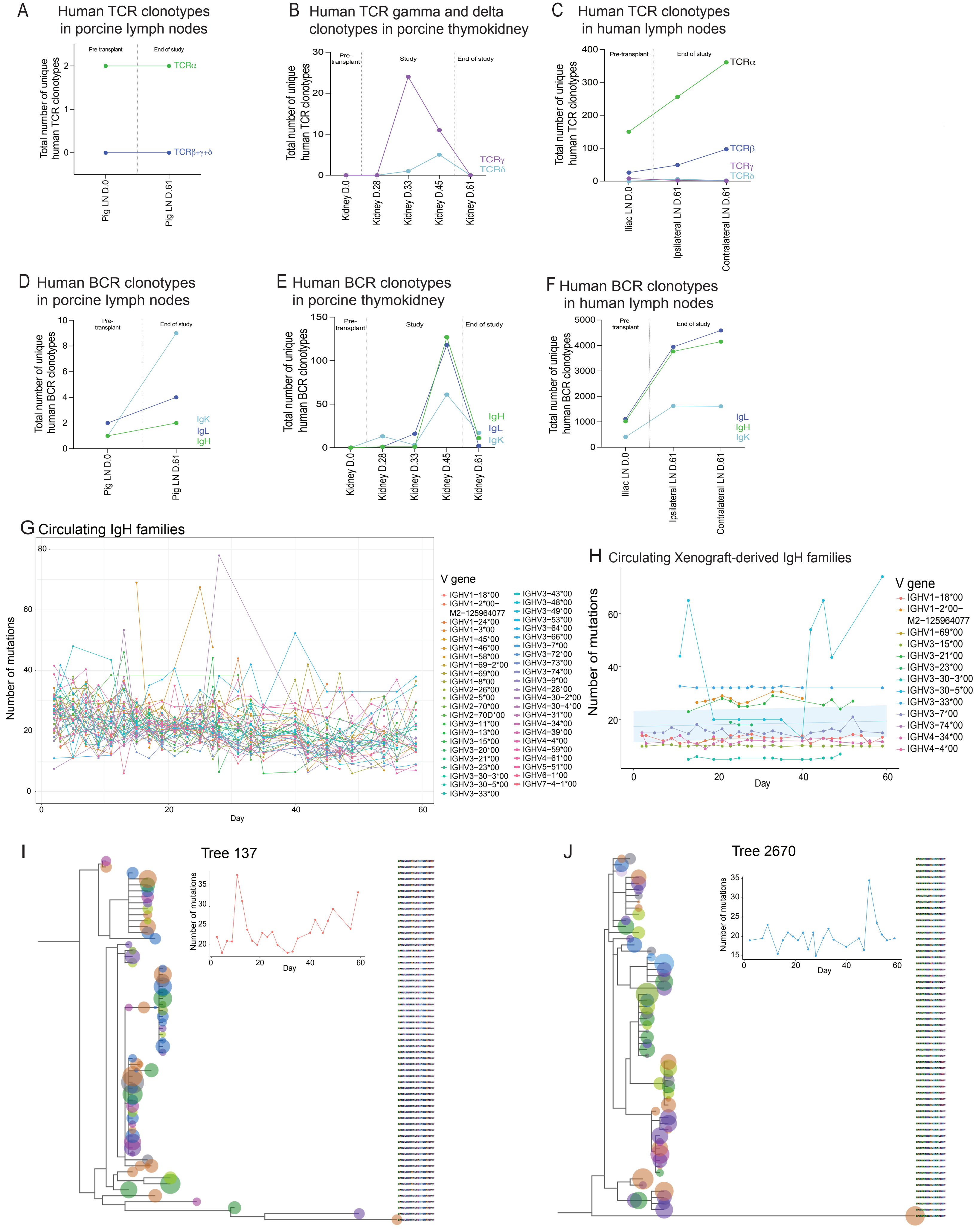

### Supplemental Figure 3

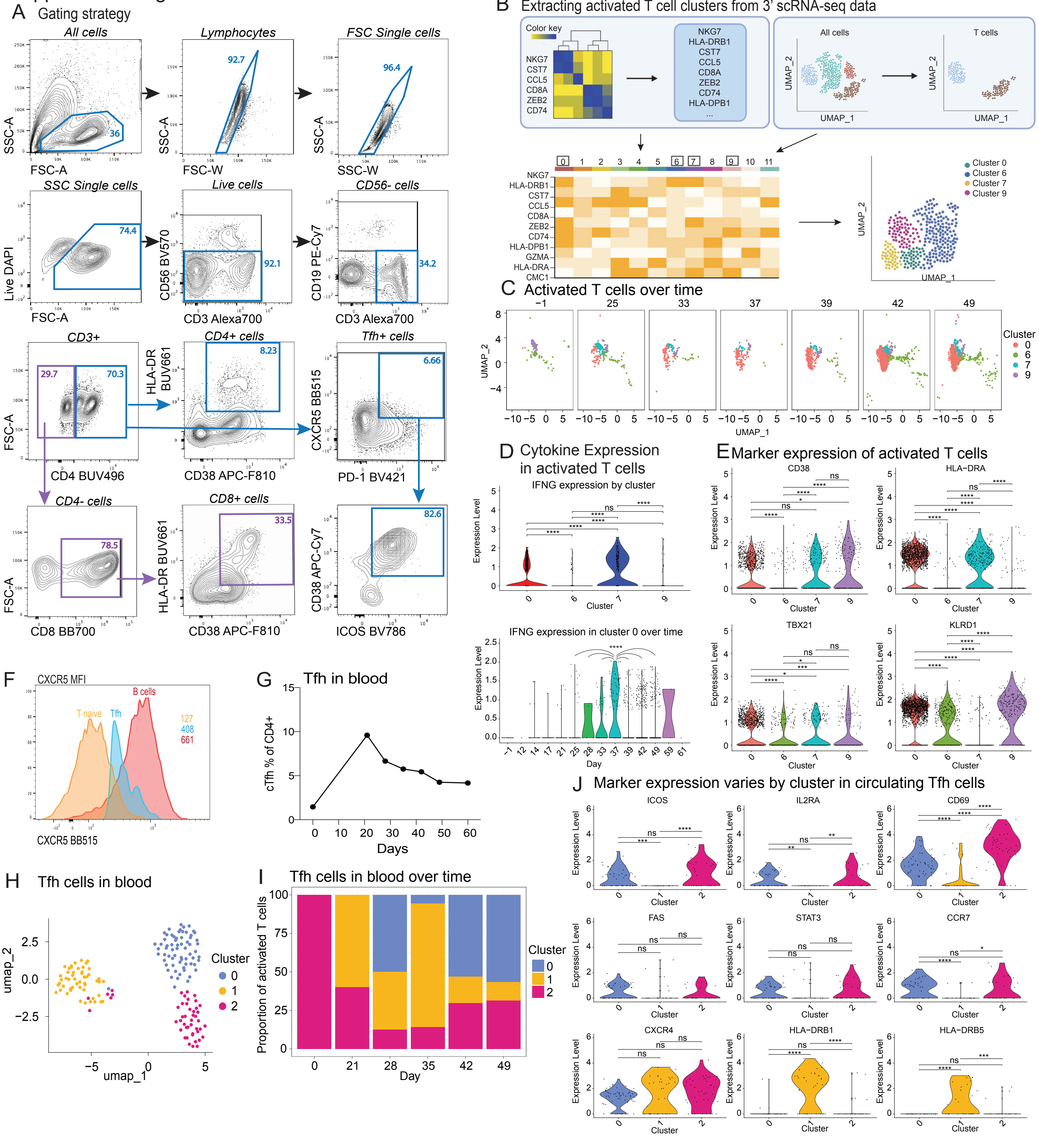

Supplemental Figure 4

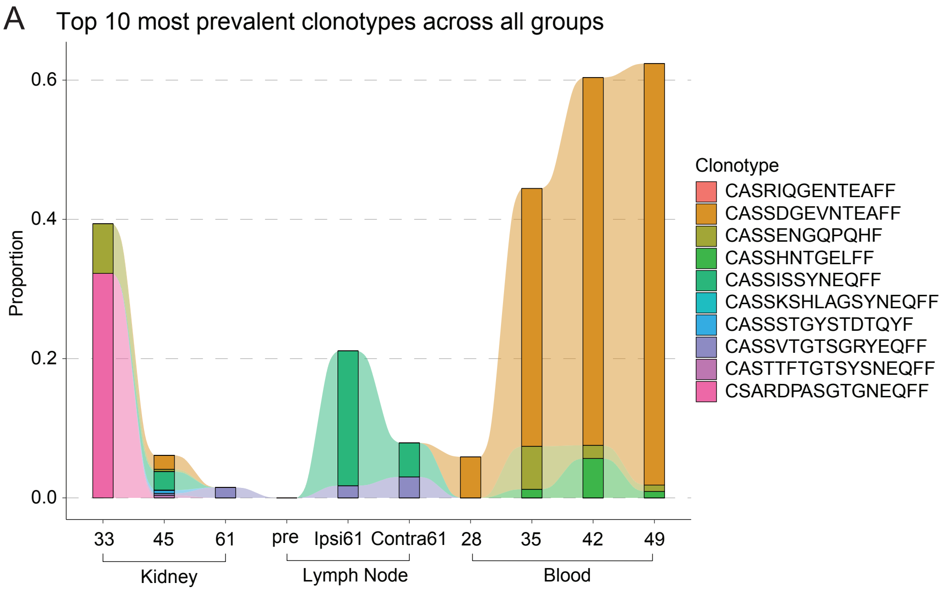


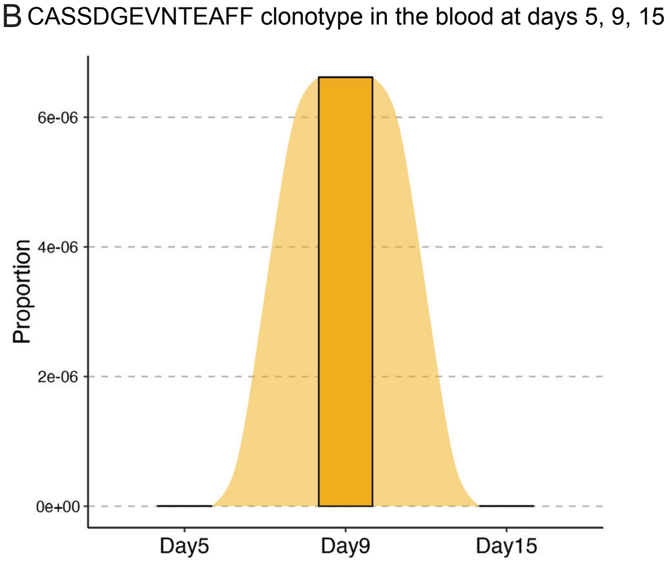

**Supplemental Table 1. RNA extraction from FFPE slides for high-throughput TCR sequencing.**

| Sample | Concentration, ng/ul | Sample volume, uL |
|---|---|---|
| Human decedent iliac lymph node prior to transplantation | 22 | 50 |
| Porcine lymph node prior to transplantation | 112.2 | 50 |
| Porcine thymokidney prior to transplantation | 160.0 | 50 |
| Porcine thymokidney, Day 28 after transplantation | 1.4 | 50 |
| Porcine thymokidney, Day 33 after transplantation | 12.0 | 50 |
| Porcine thymokidney, Day 45 after transplantation | 161.0 | 50 |
| Porcine kidney, Day 61 after transplantation, end of study | 5.1 | 50 |
| Porcine perigraft lymph node, Day 61 after transplantation, end of study | 5.7 | 50 |
| Decedent graft draining ipsilateral lymph node, Day 61 after transplantation, end of study | 18.9 | 50 |
| Decedent graft non draining contralateral lymph node, Day 61 after transplantation, end of study | 18.8 | 50 |

**Supplemental Table 2. Fluorochrome reagents and antibody clones used for FACS.**

| Antibody target | Clone | Fluorochrome | Supplier | Catalog number | RRID |
|---|---|---|---|---|---|
| CD3 | UCHT1 | Alexa700 | BD Bioscience | 557943 | AB_396952 |
| CD56 | 5.1H11 | BV570 | Biolegend | 362540 | AB_2565918 |
| CD19 | SJ25C1 | PE-Cy7 | BD Bioscience | 612938 | AB_2870221 |
| CD4 | SK3 | BUV496 | BD Bioscience | 612937 | AB_2916881 |
| CD38 | HIT2 | APC-Fire810 | Biolegend | 303549 | AB_2860783 |
| HLA-DR | G46-6 | BUV661 | BD Bioscience | 612980 | AB_2870252 |
| CXCR5 | RF8B2 | BB515 | BD Bioscience | 564624 | AB_2738871 |
| PD-1 | EH12.2H7 | BV421 | Biolegend | 329920 | AB_10960742 |
| CD8 | RPA-T8 | BB700 | BD Bioscience | 566451 | AB_2744459 |
| LiveDead | n/a | Blue Fluorescent dye | Invitrogen | 50-112-1530 | NA |
